# Systematic monitoring of 2-Cys peroxiredoxin-derived redox signals unveiled its role in attenuating carbon assimilation rate

**DOI:** 10.1101/2021.02.03.429619

**Authors:** Nardy Lampl, Idan Nissan, Raz Lev, Gal Gilad, Matanel Hipsch, Shilo Rosenwasser

## Abstract

Transmission of reductive cues from the photosynthetic electron transport chain to redox-regulated proteins plays a crucial role in activating chloroplast metabolism. However, deciphering the role of their counterbalanced oxidative signals is challenging due to monitoring difficulties. Here, we demonstrate the light-depended redox modification of chloroplast-targeted 2-Cys peroxiredoxins and introduce peroxiredoxin-based biosensors to monitor photosynthetically-derived oxidative signals. By employing a set of genetically encoded biosensors, we show the induction of oxidative signals under habitual light intensities and their inverse relationship with NADPH levels, unraveling the combined activity of reducing and oxidizing signals in fine-tuning chloroplast metabolism. A faster increase in carbon assimilation rates during photosynthesis induction phase was measured in plants deficient in 2-Cys peroxiredoxins compared to wild-type, suggesting the involvement of oxidative signals in attenuating photosynthesis under variable light environments. We suggest that oxidative signals measured by peroxiredoxin-based biosensors reflect the limitation to photosynthesis imposed by the redox regulatory system.

**One-Sentence Summary:** A genetically encoded biosensor unmasked the dominant role of photosynthetically-derived oxidative signals under habitual conditions.

## Introduction

Sophisticated mechanisms enabling adjustments of light-capturing reactions and downstream metabolic processes have evolved in sessile plants, allowing them to reach optimal photosynthetic performance. Under low-light conditions, efficient photosynthetic activity and tight regulation of the reducing power distribution between essential metabolic pathways are required for optimal utilization of the available energy input. On the other hand, energy dissipation under high-light (HL) conditions is critical to avoid over-production of harmful reactive oxygen species (ROS)^1^. Reductive and oxidative signals, both emanating from the photosynthetic electron transport chain (PETC), and their transfer to regulated thiol proteins, play a significant role in linking the performance of photosynthesis to chloroplast transcription, translation, and metabolic processes, thereby enabling rapid acclimation to instantaneous changes in daytime photon fluxes^2–5^.

Reductive signals in chloroplasts are mainly generated by the flux of electrons derived from ferredoxin to target proteins, via ferredoxin-dependent thioredoxin reductase (FTR), to thioredoxins (Trxs)^6^ and via NADPH-dependent thioredoxin reductases C (NTRC)^7^, thereby linking chloroplast metabolism to light availability and photosynthetic activity. Counteracting oxidative signals are mainly generated from electron flux through the water-water cycle (WWC). During the WWC, electrons are donated from photosystem I (PSI) to molecular oxygen in the Mehler reaction, yielding superoxide radicals (O_2_^·−^), which are then dismutated to molecular oxygen (O_2_) and H_2_O_2_ in a reaction catalyzed by superoxide dismutase (SOD)^8–10^. The production of H_2_O_2_ through the WWC allows communication between the photosynthetic light reactions and downstream metabolic processes by transmitting oxidative signals to redox-regulated proteins^11–14^

The detoxification of photosynthetically produced H_2_O_2_ is mediated by chloroplast-targeted ascorbate peroxidases (APXs) and 2-Cys peroxiredoxins (2-Cys Prxs)^15,16^. The latter are highly abundant and remarkably efficient thiol peroxidases, displaying high catalytic efficiency of up to 10^8^ M^−2^ s^−1 17,18^. Electrons delivered from the PETC, reduce 2-Cys Prx via the FTR/TRX pathways or through NADPH via NTRC activity^19–22^. Therefore, the 2-Cys Prxs redox state is determined by the balance between photosynthetically produced reducing and oxidizing equivalents. Importantly, recent studies have demonstrated oxidative signal transmission, mediated by 2-Cys Prxs, from H_2_O_2_ to target proteins at the onset of the light and dark periods and under low- and fluctuating-light conditions^23–27^. Despite the considerable importance of the regulatory role of 2-Cys Prx, most of our current knowledge is based on measuring its oxidation state using western blot analysis or examination of mutant lines performance, rendering it challenging to detect the transmission of oxidative signals in real-time and with high temporal resolution.

Genetically encoded redox-sensitive green fluorescent protein (roGFP) probes enable, due to their high sensitivity and reversibility, systematic *in vivo* mapping of H_2_O_2_ production and the glutathione redox state (*E_GSH_*), in different sub-cellular compartments^28–35^. In vitro characterization of roGFP showed that its reduction is mediated by Glutaredoxins (Grxs), which catalyze the reversible electron flow between GSH and target proteins ^31,32^. Recently, an H_2_O_2_ sensor based on a redox relay mechanism between Tsa2, a highly efficient *S. cerevisiae* 2-Cys peroxiredoxin, and roGFP2 was developed^36^. To increase the probe’s sensitivity to low H_2_O_2_ levels, a mutation in the resolving cysteine (C_R_) of Tsa2 (roGFP2-Tsa2ΔC_R_) was introduced, thus preventing competition between the Trx-mediated reduction of the Tsa2 moiety and roGFP oxidation. The resulting probe enabled the measurement of endogenous basal H_2_O_2_ levels in yeast and *Chlamydomonas reinhardtii* cells^36,37^; however, such a roGFP2 Prx-based probe has not been yet implemented to measure the dynamic oxidation of 2-Cys Prx in higher plants.

Here, we employed chl-roGFP2-PrxΔCR, a new 2-Cys Prx biosensor, to establish the role of 2-Cys Prx in light-dependent oxidative signal transmission in *Arabidopsis* plants. By parallel measurements of the endogenous 2-Cys Prx and chl-roGFP2-PrxΔC_R_ oxidation states, we showed that 2-Cys Prx is directly regulated by light intensity. Furthermore, the dynamics of 2-Cys Prx-dependent oxidative signals, throughout the day, under various light conditions, were resolved by systematic monitoring of the probe’s redox state and showed a negative correlation with the availability of chloroplast NADPH. Finally, by using a set of genetically encoded biosensors, we unveil the combined effect of the reductive and oxidative pathways in fine-tuning photosynthetic activity.

## Results

We sought to explore the direct effect of light intensity on the redox state of 2-Cys Prx. In agreement with a previous report^23^, transferring plants from dark to normal growth light (GL, 120 μmol photons m^−2^ s^−1^) at the beginning of daylight, resulted in a reduction of 2-Cys Prx catalytic sites, as reflected by the dominance of the dimer (oxidized) band under dark (at the end of the night) and by the prolonged decrease in its relative abundance under light conditions (Fig. 1A). Importantly, this reduction was not observed in plants that were pretreated with 3-(3,4-dichlorophenyl)-1,1-dimethylurea (DCMU), an inhibitor of photosynthetic electron transport (Fig. 1A), simulating dark conditions. To distinguish between the fully oxidized (2S-S) and half-oxidized (1S-S) forms, a gel-shift assay was carried out with methoxypolyethylene glycol-maleimide (mPEG). A mixture of three bands was identified, with the higher and middle bands trapped with mPEG and corresponding to the 2S-S and 1S-S forms, respectively (Fig. 1B, C). Under dark conditions, a relatively high level of the fully oxidized form was visible. On transfer of plants from dark to GL, the relative levels of the fully reduced form significantly increased and were paralleled by a decrease in the fully oxidized form. Such a reduction was not detected in plants treated with DCMU under GL, indicating that PETC was the source of the reduction power.

**Fig. 1.**
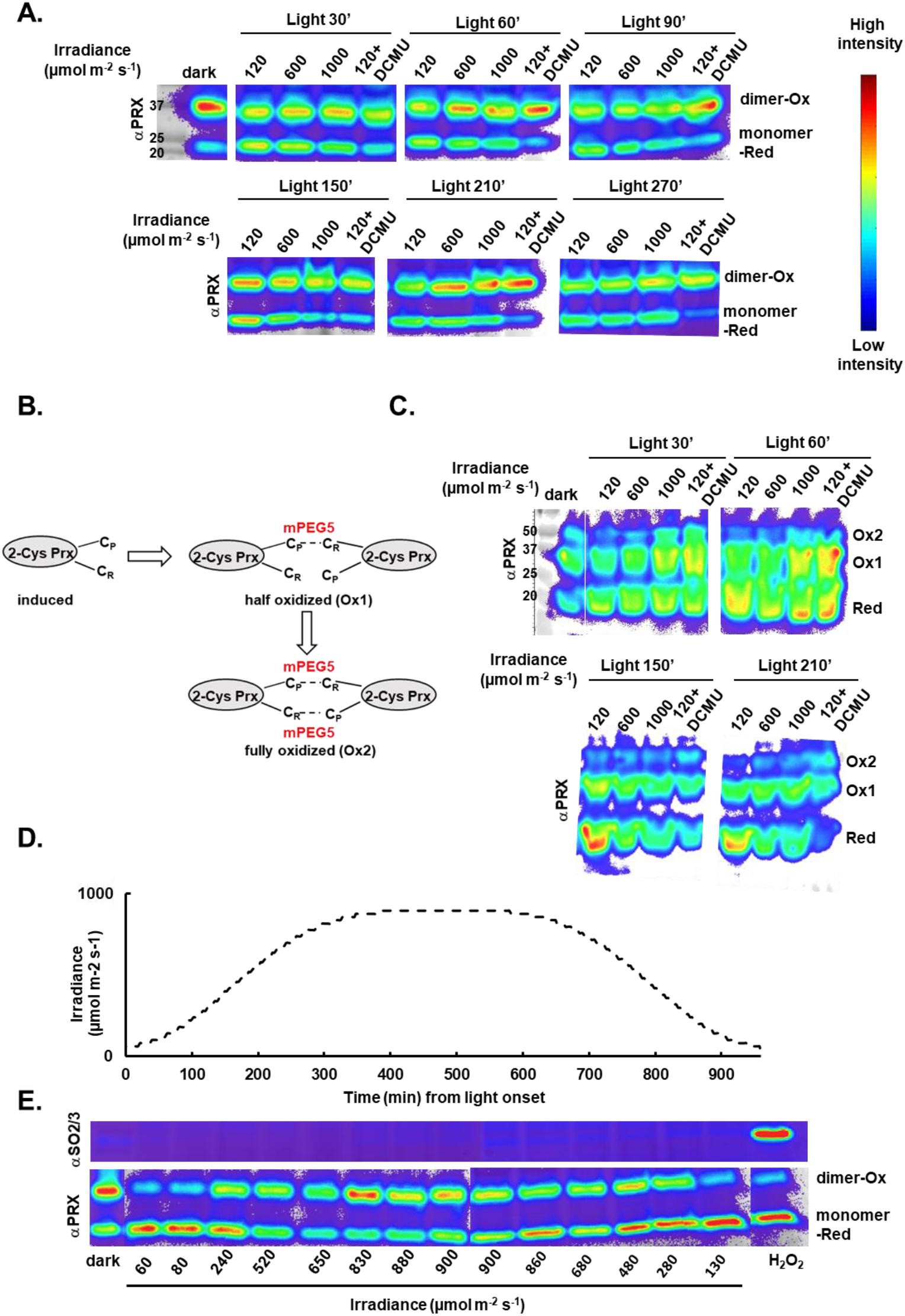
Light responsiveness of the 2-Cys Prx redox state. Plants were kept in the dark for 8 h and then exposed for 5.5 h to the indicated light intensities. Total protein was extracted using trichloroacetic acid (TCA), and free thiols were blocked with N-ethylmaleimide (NEM). (**A**) An immunoblot, using anti-Prx, showing the oxidized and reduced states of 2-Cys Prx, as dimers and monomers, respectively. (**B**) Schematic illustration of a redox mobility gel assay (with samples from A). (**C**) Gel-shift assay with methoxypolyethylene glycol-maleimide (mPEG) showing the in vivo redox state of 2-Cys Prx residues in plants exposed to light intensities as in A. Following NEM treatment, 1,4-Dithiothreitol (DTT) was applied to reduce disulfides before mPEG labeling. The number of oxidized Cys residues (Ox1 and Ox2) was determined by the increased molecular mass of 2-Cys Prx after labeling with mPEG. (**D**) Light intensity over time. (**E**) Plants were exposed to the light regime indicated in D. Immunoblot assay showing the oxidized and reduced states of 2-Cys Prx, as dimers and monomers, respectively (anti-Prx) and 2-Cys Prx hyperoxidation (anti-SO_3_).

Plant exposure to high light (HL) intensities (600 or 1000 μmol photons m^−2^ s^−1^) (Fig. 1A) resulted in a similar reduction of 2-Cys Prx upon dark-to-light transition for the two tested HL intensities, which stabilized at a higher oxidation state as compared with GL. A significant amount of fully oxidized 2-Cys Prx arose in plants exposed to 1000 μmol photons m^−2^ s^−1^ (Fig. 1B, C), suggesting that increased illumination enhanced 2-Cys Prx dimer formation, in which both catalytic sites participate in disulfide exchange reactions. Plant exposure to constant HL conditions or DCMU also led to higher levels of hyperoxidized 2-Cys Prx than GL, as measured using SO_3_ antibodies (Supplementary Fig. 1).

An apparent light-dependent oxidation response of 2-Cys Prx was observed under prolonged, physiologically relevant light conditions (Fig. 1D, E). Interestingly, the major shift in oxidation was during the non-stressed light regime, in the transition from low (80-130 μmol photons m^−2^ s^−1^) to moderate (240-280 μmol photons m^−2^ s^−1^) light intensities, at the initiation and termination of the light phase (Fig. 1D, E), implying its signaling role under habitual light conditions. Notably, accumulation of the Prx hyperoxidation form was not detected (Fig. 1E), suggesting that a gradual increase in light conditions inhibits 2-Cys Prx overoxidation even while reaching HL conditions. These observations demonstrated that the 2-Cys Prx redox state is directly affected by the PETC and that its oxidation state under gradual changes in light intensities that mimic natural conditions in the field, is firmly linked to changes in light intensities.

We recognized that the observed oxidation patterns of 2-Cys Prx in response to changing light intensities reflect the balance between its peroxidase activity and its reduction by TRXs. Therefore, to explicitly inspect the oxidative signals mediated by 2-Cys Prx activity, we generated, based on a previous work in yeast ^36^, a chl-roGFP2-PrxΔC_R_ probe by genetically fusing roGFP2 to a mutated version of 2-Cys Prx A (BAS1), in which the resolving cysteine (C_R_) was changed to alanine, impeding its TRX-dependent reduction (Fig. 2A-C). BAS1 was fused at its N terminus to roGFP2, since initial attempts to fuse it at its C terminus resulted in probe cleavage (Supplementary Fig. 2). Interestingly, the cleaved roGFP obtained from the fusion of BAS1 at its C-terminus was localized in the chloroplast (Supplementary Fig. 2), implying that the C terminus of BAS1 is prone to cleavage by specific chloroplast proteases. A dynamic range of 5.6 was calculated between the fully oxidized (R_oxi_=0.45) and fully reduced (R_red_=0.08) forms of chl-roGFP2-PrxΔCR, ensuring a sufficient signal to noise ratio (Fig. 2D). Whole-plant ratiometric analysis showed a higher oxidation state of the chl-roGFP2-PrxΔC_R_ probe compared to chl-roGFP2 under steady-state conditions (GL) (~ 60% and ~ 40%, respectively, Fig. 2E-G), a phenomenon also observed with the roGFP2-Tsa2ΔC_R_ probe^36,37^, indicating a direct effect of 2-Cys Prx on the roGFP moiety. Shifting plants from dark to GL induced a slight change in the redox state of chl-roGFP2-PrxΔC_R_ and significant changes were visualized in its oxidation degree (OxD) upon shifting plants from dark to different HL intensities (450, 750, or 1000 μmol photons m^−2^ s^−1^), reaching its almost fully oxidized state within 5 min of transition to light (Fig. 2E, G). In contrast, the unfused chl-roGFP2 probe was only moderately oxidized under HL conditions (Fig. 2E, F). These findings demonstrate the enhanced light sensitivity of chl-roGFP2-PrxΔC_R_ compared to chl-roGFP2, achieved by positioning BAS1 in close proximity to roGFP2. As both probes are subjected to reduction in a GSH/GRX-dependent manner, the comparison between their dynamic oxidation enables resolution of the effect on H_2_O_2_-dependent thiol oxidases mediated by BAS1.

**Fig. 2.**
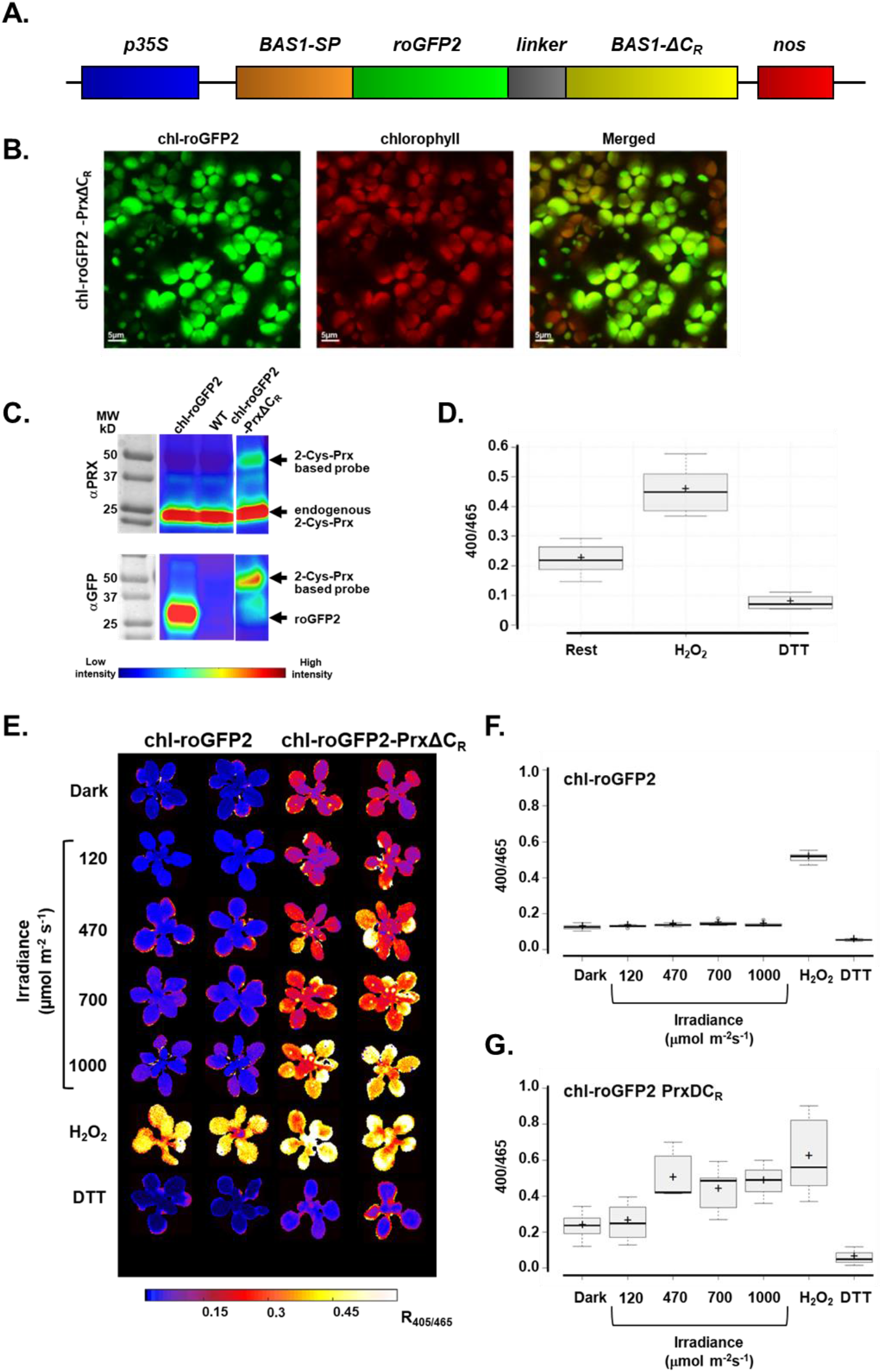
Light responsiveness of chloroplast-targeted peroxiredoxin-based redox probes. (**A**) Schematic diagram of the gene cassette used to transform *Arabidopsis* plants. (**B**) Subcellular localization of chl-roGFP2-PrxΔC_R_ in mesophyll cells, as detected by confocal microscopy. (**C**) Detection of the fused sensor protein by western blot analysis using either anti-Prx or anti-GFP antibodies. (**D**) Fluorescence ratios (405/465) in plants expressing chl-roGFP2-PrxΔC_R_ under steady-state conditions and following treatments with 100 mM DTT or 500 mM H_2_O_2_. Fluorescence emitted from three-week-old plants grown in soil was recorded using a plate reader. (**E**) Whole-plant ratiometric analysis of chl-roGFP2 and chl-roGFP2-PrxΔC_R_ fluorescence under dark and 5 min following transfer from dark to different light intensities. Ratiometric analysis of plants treated with 100 mM DTT or 500 mM H_2_O_2_ is also presented. The roGFP2 fluorescence images were captured at 510 nm, following excitation at 400 nm and 465 nm. (**F, G**) Quantification of ratiometric images, presented as box-plots.

We further investigated the daily dynamics of the chl-roGFP2-PrxΔC_R_ oxidation state in comparison to chl-roGFP2 using an automated system that allows for continuous measurements of fluorescence signals emanating from living plants expressing genetically encoded biosensors (Fig. 3A-C, Methods). Monitoring probe oxidation throughout the day further demonstrated the higher sensitivity of chl-roGFP2-PrxΔC_R_ to light compared to chl-roGFP2. While both probes were oxidized upon transition from dark to light under all examined light conditions, significantly higher oxidation values were recorded under HL in chl-roGFP2-PrxΔC_R_ (~85-95%) as compared to chl-roGFP2 plants (~65-70 %; Fig. 3B-C).

**Fig. 3.**
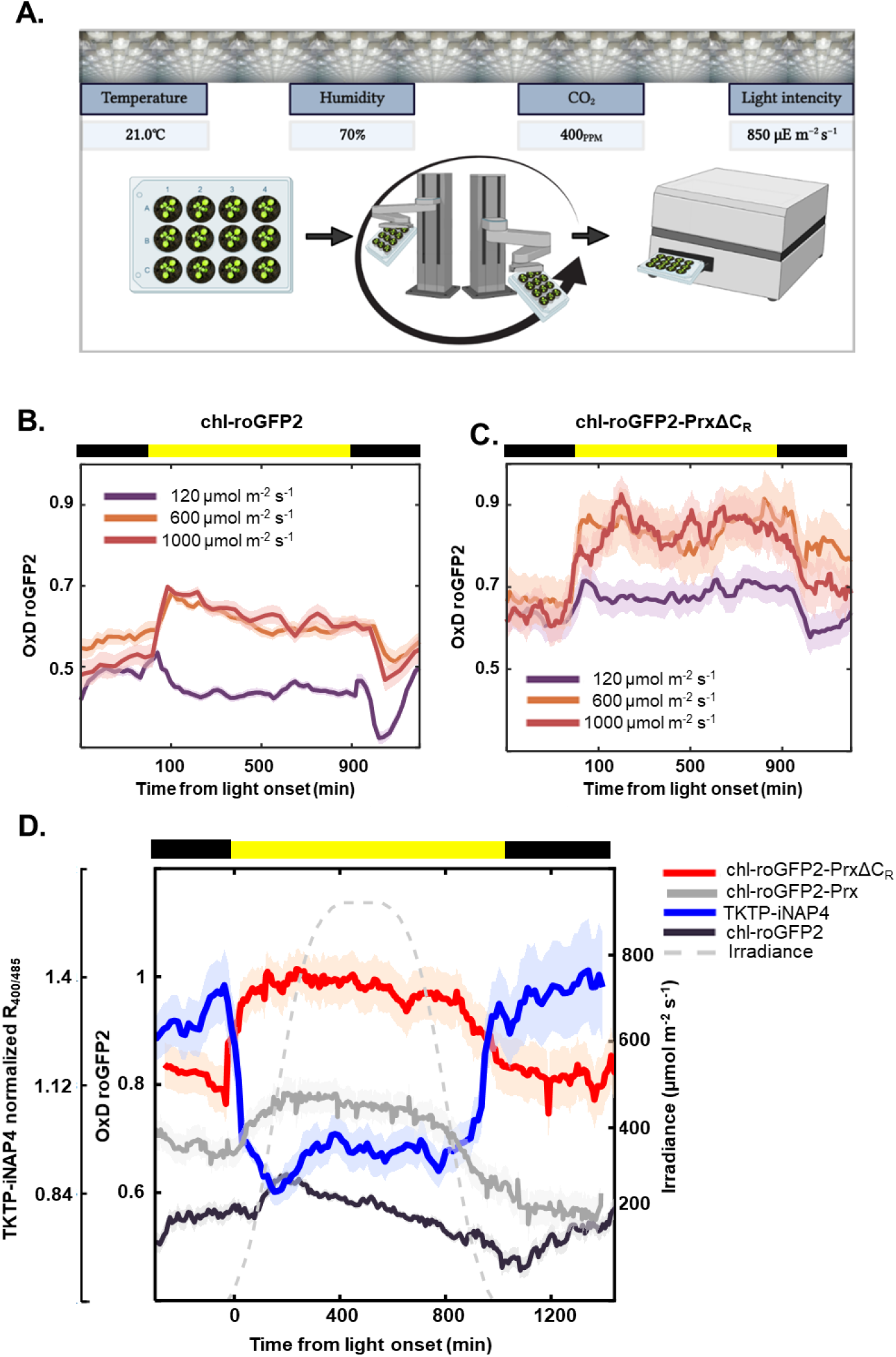
Systematic monitoring of light-induced oxidative signals. (**A**) Schematic illustration of an automated system used to monitor fluorescence signals in living plants. Illustration partially created with BioRender.com. (**B, C**) Dynamic changes in chl-roGFP2 and chl-roGFP2-PrxΔC_R_ oxidation degree (OxD) in plants exposed to growth light (GL) (120 μmol photons m^−2^ s^−1^) or high light (HL) (600 or 1000 μmol photons m m^−2^ s^−1^) after 8 h in the dark. (**D**) Dynamic changes in OxD of chl-roGFP2-PrxΔC_R_, chl-roGFP2-Prx and chl-roGFP2 as well as TKTP-iNAP4 ratio (400/485nm) are presented throughout the day. The applied light intensities are depicted as a dashed yellow line. For each line, between 24 and 40 plants divided into 3-5 independent plates were analyzed and consolidated in a “sliding window” (n=3-5) display. Values represent means ± SE. The color bar denotes the light conditions: black-dark, yellow-light.

To characterize chl-roGFP2-PrxΔC_R_ oxidation dynamics under a range of light intensities, we monitored their OxD in plants exposed to gradual changes in light intensities that mimic natural conditions, as described in Fig. 1E. For comparison, the oxidation dynamics of the chl-roGFP2 and chl-roGFP2-Prx probes (without the mutation in the C_R_, Supplementary Fig. 4), were also monitored, as was NADPH availability, using the recently developed *Arabidopsis* NADPH sensor (TKTP-iNAP4)^38,39^. As shown in Fig. 3D, the four probes exhibited entirely different patterns. The chl-roGFP2-PrxΔC_R_ OxD sharply increased from the light onset, reaching its full oxidation state (~100%) after approximately two hours, when the light intensity was ~300 μmol photons m^−2^ s^−1^. A gradual reduction in OxD was only detected 14 h after light onset, when light intensities were dimmed again to ~300 μmol photons m^−2^ s^−1^. We recorded a lower magnitude of oxidation of chl-roGFP2-Prx compared to chl-roGFP2-PrxΔC_R_, supporting the higher sensitivity of the latter to oxidative signals, achieved by the C_R_ mutation. Notably, the oxidative patterns of chl-roGFP2-PrxΔC_R_ and chl-roGFP2-Prx probes in response to changes in light intensities were similar to those of endogenous 2-Cys Prx (Fig. 3D and Fig. 1E), reinforcing the probe’s capabilities to detect 2- Cys Prx-mediated redox signals. We reasoned that the differences between chl-roGFP2-PrxΔC_R_ and chl-roGFP2-Prx OxD reflected reducing equivalents transferred to 2-Cys Prx by the combined activity of NTRC and TRXs. In contrast to the oxidation patterns of chl-roGFP2-Prx and chl-roGFP2-PrxΔCR, the unfused chl-roGFP2 showed a gradual increase in OxD, reaching its highest level (65%) after 3 h, when the light intensity was ~600 μmol photons m^−2^ s^−1^. From this point onward, a gradual decrease in chl-roGFP2 oxidation was observed, reaching its lowest OxD (42%) at the beginning of the dark period.

Strikingly, the redox state of chl-roGFP2-PrxΔC_R_ negatively correlated with the dynamic changes in stromal NADPH availability, as recorded using the TKTP-iNAP4 lines. While the NADPH level was high and stable at night, a sharp decrease was detected under light, starting after light onset and reaching its lowest level at ~300 μmol photons m^−2^ s^−1^. A gradual increase in NADPH level was observed ~14 h after light onset, when light intensities were lowered, until it reached its highest level at the beginning of the night. These results imply that the availability of stromal NADPH governs the induction of photosynthetically-derived oxidative signals.

The significant oxidation of chl-roGFP2-PrxΔC_R_ under low to moderate light intensities (Fig.3D), which matches the increasing recognition that 2-Cys Prx plays a regulatory role under low-light conditions^23–25,40^ drove us to further investigate the sensitivity of the chl-roGFP2-PrxΔC_R_ redox state to low-light intensity. To this end, plants were exposed to a gradual increase in light intensities, starting with ~22 μmol photons m^−2^ s^−1^ and increasing up to 330μmol photons m^−2^ s^−1^ (Fig. 4A). An immediate reduction of chl-roGFP2 was observed at ~22 μmol photons m^−2^ s^−1^, followed by oxidation at a light intensity of ~44 μmol photons m^−2^ s^−1^, with oxidation reaching a maximum under 330 μmol photons m^−2^ s^−1^. In contrast, gradual oxidation of chl-roGFP-PrxΔC_R_ was observed with progression through the low-light gradient. Stromal NADPH availability negatively correlated with chl-roGFP2-PrxΔC_R_ as in Fig. 3D. To further examine the induction of reductive and oxidative signals under low light, we monitor the probe’s oxidation pattern during prolonged exposure to low light. Interestingly, the chl-roGFP2-PrxΔC_R_ and chl-roGFP2-Prx probes showed contrasting responses upon transition from dark to low light (Fig. 4B). While chl-roGFP2-PrxΔC_R_ OxD slightly increased when plants were transferred to low light, chl-roGFP2-Prx underwent a significant reduction. This reduction continued for at least two hours from light onset until stabilization at a lower OxD (35%). The differences between the OxD of roGFP2-PrxΔCR, chl-roGFP2-Prx, and the unfused chl-roGFP2 after the onset of low light demonstrate the high magnitude of reductive signals under low-light intensities, emanating from TRXs and GSH/GRXs activity and which are attenuated by induced oxidative signals (Fig. 4C).

**Fig. 4.**
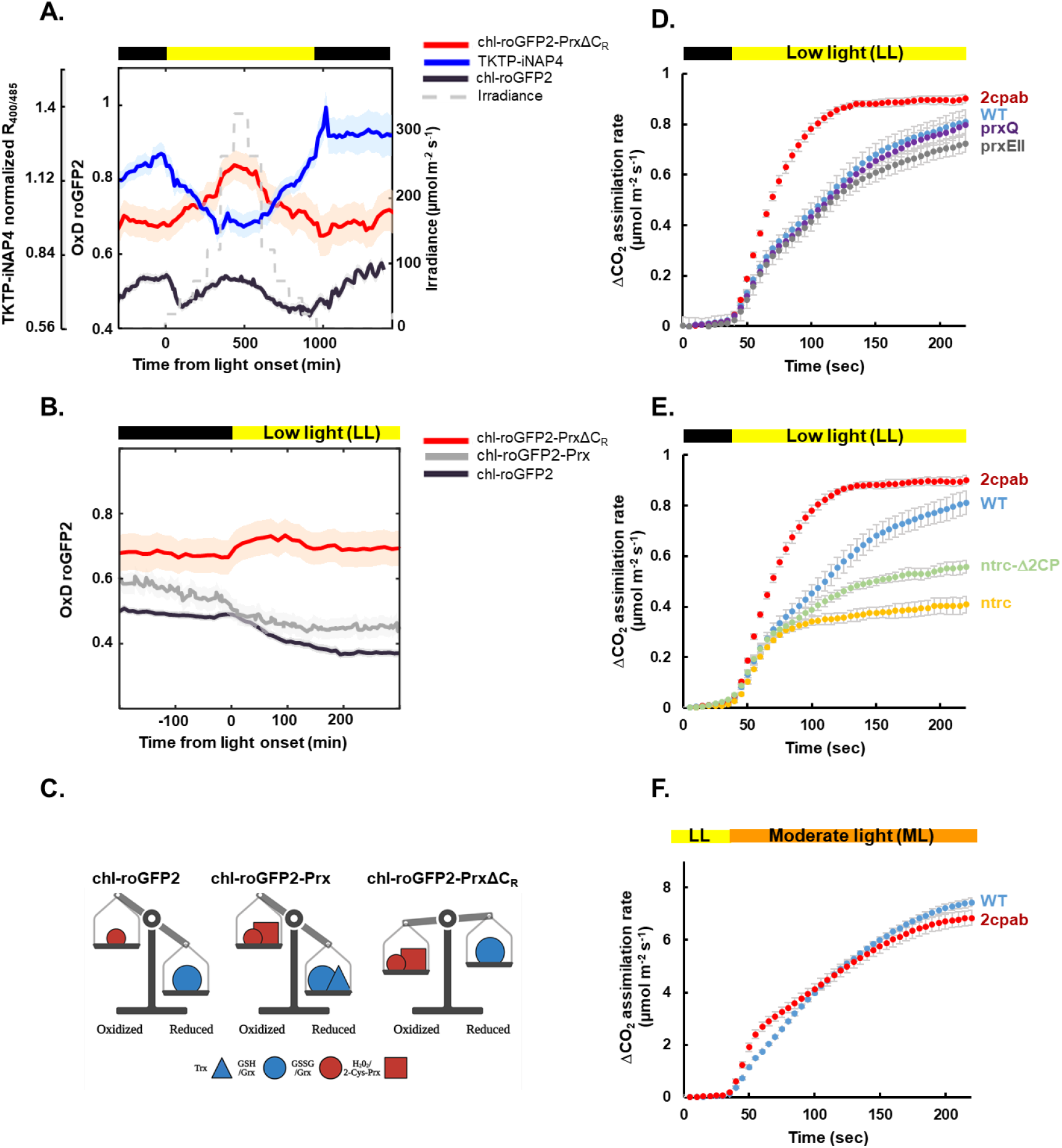
chl-roGFP2-PrxΔC_R_ reports on the induction of oxidative signals under low light. (**A**) Daily dynamic changes in the oxidation degree (OxD) of chl-roGFP2-PrxΔC_R_ and chl-roGFP2, and in TKTP-iNAP4 ratios, in plants exposed to a gradual increase (low to moderate) light intensities followed by an equivalent decrease, are presented. Monitoring was performed in 3-week-old plants grown in 12-well plates. For each line, between 24 and 40 plants divided into 3-5 independent plates were analyzed and consolidated in a “sliding window” (n=3-5) display. Values represent means ± SE. The color bar denotes the light conditions: black-dark, yellow-light. (**B**) Dynamic changes in chl-roGFP2-PrxΔC_R_, chl-roGFP2-Prx and chl-roGFP2 OxD at the onset of low light (22 μmol photons m^−2^ s^−1^). The applied light intensities are depicted as a dashed yellow line. (**C**) A proposed model of the oxidation and reduction powers that shape the oxidation state of the chl-roGFP2-PrxΔC_R_, chl-roGFP2-Prx and chl-roGFP2 sensors. Figure created with BioRender.com. (**D, E**) Carbon assimilation rate was measured in dark-adapted WT and mutants plants (*2cpab, ntrc, ntrc-Δ2CP, prxQ* and *prxEII*) upon exposure to low light (22 μmol photons m^−2^ s^−1^). As differences in dark respiration rates were observed between the mutant lines, gas exchange data were normalized to the dark values. Values for the carbon assimilation are presented as means of 4-5 pots with 4 plants in each ± SE. The color bar denotes the light conditions: black-dark, yellow-light. (**F**) Carbon assimilation rate was measured in moderate light (330 μmol photons m^−2^ s^−1^) adapted WT and *2cpab* plants upon exposure to 5min of low light (22 μmol photons m^−2^ s^−1^) followed by moderate light (330 μmol photons m^−2^ s^−1^). Gas exchange data were normalized to the dark values. Values for the carbon assimilation are presented as means of 4 pots with 4 plants in each ± SE. The color bar denotes the light conditions: yellow-low light, orange-moderate light.

We further hypothesized that the observed oxidative signals, upon transition from dark to low light, play a role in Calvin–Benson cycle (CBC) enzymes deactivation, thus substantially influencing carbon assimilation rates. To assess to what extent 2-Cys Prxs limit photosynthesis, we measured the increase in carbon assimilation rates during the photosynthesis induction phase in dark-adapted mutants deficient in 2-Cys Prxs (*2cpab*)^26^ and wild type (WT) plants. Notably, while upon transition to low light, a considerable lag in the attainment of photosynthesis steady-state was observed in WT plants, a significantly faster induction of photosynthesis was detected in *2cpab* plants (Fig. 4D, Supplementary Fig. 5), pointing to the inhibitory effect of 2-Cys Prxs activity during the photosynthesis induction phase. Furthermore, this fast induction was not detected in plant lines mutated in prxQ and prxIIE, two additional chloroplast-targeted Prxs^41,42^ (Fig. 4D), suggesting the exclusive role of 2-Cys Prxs in determining the limitation to photosynthesis during its induction phase. The limitation to photosynthesis mediated by 2-Cys Prx was also demonstrated in plants mutated in NTRC, an essential component of the reductive signal pathway (Fig. 4E). As shown in Fig. 4E, *ntrc* plants showed a significantly lower increase in carbon assimilation rate than WT upon shifting of plants from dark to low light, emphasizing the dependency of the photosynthesis induction phase on the redox regulatory network. Notably, this effect was partially recovered in *ntrc-Δ2cp* triple mutant plants^43^, which express severely low levels of 2-Cys Prxs (Fig. 4E). These results are in agreement with the reported suppression of *ntrc* phenotype by decreased levels of 2-Cys Prxs^43^.

The fast induction of photosynthesis in *2cpab* compared to WT plants was also observed upon returning of moderate-light-adapted plants from short exposure of low light (5 min) back to moderate light intensities (Fig. 4F). In contrast, when *2cpab* and WT plants were exposed to a brief low light period of 1min (Supplementary Fig. 6), no differences were observed, indicating a time-dependent built-up of the 2-Cys Prx inhibitory effect. These results demonstrate the role of the redox regulatory system in shaping carbon assimilation rates under dynamic light conditions and the limitation to photosynthesis imposed by 2-Cys Prxs activity.

## Discussion

It had been established that photosynthesis is dependent on incoming light not only as an energy source to drive electron transfer but also for reducing regulatory disulfides in stromal proteins, allowing the adjustment between chloroplast metabolism and photosynthetic activity^6^. Besides generating reductive fluxes, photosynthetic light reactions also produce oxidizing agents such as hydrogen peroxide, chiefly through the water-water cycle^9^. Accordingly, the *in-vivo* redox state of regulatory thiols is dependent on the sum of reductive and oxidizing signals derived from the photosynthetic reducing and oxidizing activity, respectively (Fig. 5). While classical redox biology approaches allow the capturing of the redox state of a given protein/thiol, discerning the biological context of the reductive and oxidative pathways is still challenging. On the basis of the increasing recognition that chloroplast-targeted 2-Cys Prxs is capable of transferring oxidative signals to target proteins^23–25,27,43^ and based on the recently developed roGFP2-Tsa2ΔC probe in yeast^36^, we developed chl-roGFP2-PrxΔC_R_ in order to define the exact light regime and time frame in which oxidizing signals are transmitted.

**Fig. 5:**
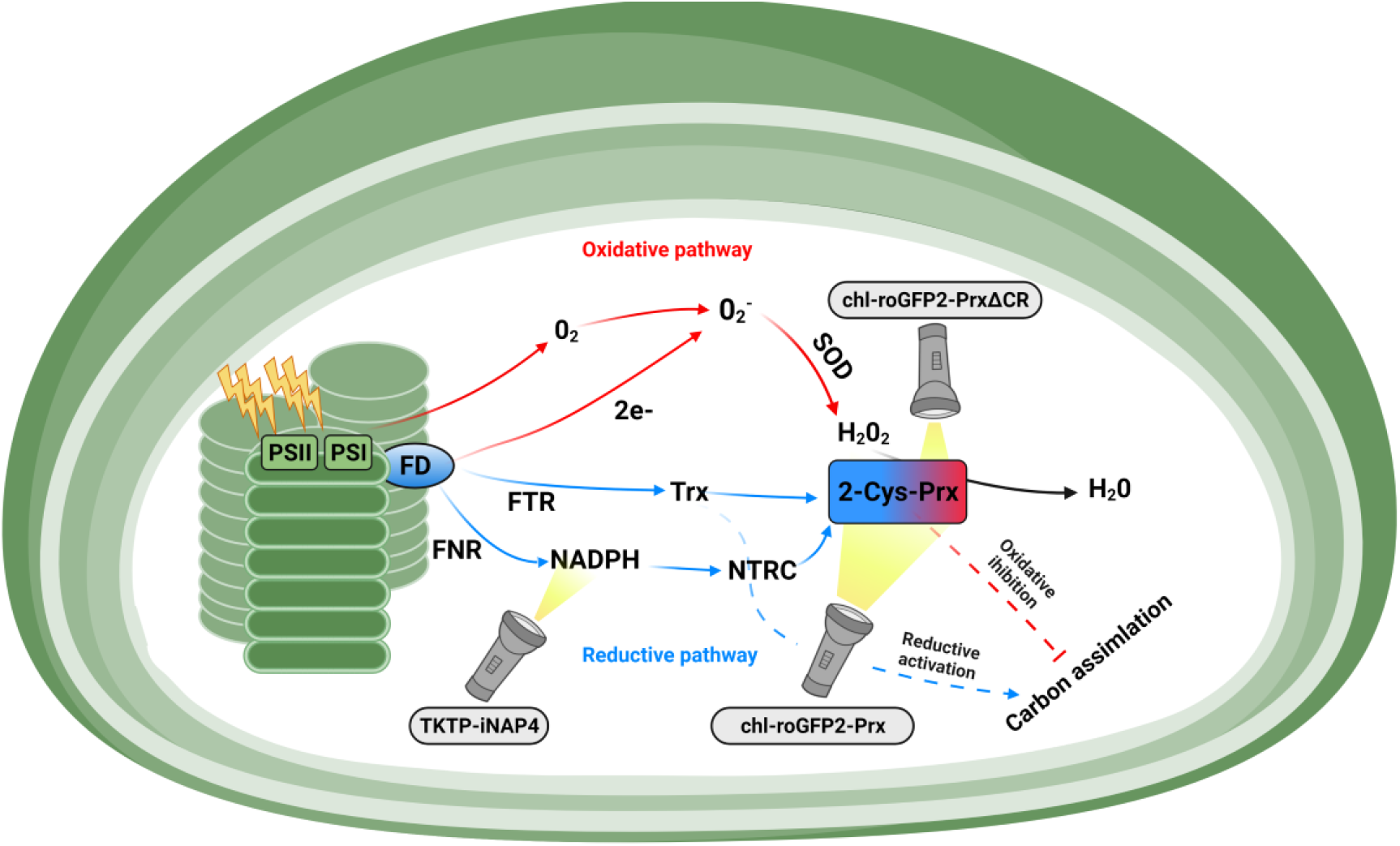
A proposed schematic model of redox signals emanating from the photosynthetic electron transport chain and their detection using genetically encoded biosensors. Reductive signals are generated by the flux of electrons derived from ferredoxin to target proteins, via ferredoxin-dependent thioredoxin reductase (FTR), to thioredoxins (Trxs) and via ferredoxin-NADP(+) reductase (FNR) and NADPH-dependent thioredoxin reductases C (NTRC). Counterbalanced oxidative signals are generated from electron flux through the water-water cycle via the activity of 2-Cys Prxs. Both signals instantaneously fine-tune the redox state of CBC enzymes and consequently carbon assimilation rates. While the redox state of the chl-roGFP2-Prx sensors is shaped by the sum of the reductive and oxidizing powers working on 2-Cys Prxs, the chl-roGFP2-PrxΔC_R_ sensors measure the oxidative signals exclusively. TKTP-iNAP4 is used to assess NADPH availability in chloroplasts. Figure created with BioRender.com.

Interestingly, by using a set of genetically encoded biosensors (Fig. 5), we demonstrated that oxidizing activity governed by 2-Cys Prxs is prevalent under habitual growth conditions. This postulation is based on the differences in oxidation patterns observed for the chl-roGFP2-PrxΔCR, and two additional probes monitored in this study, chl-roGFP2-Prx and chl-roGFP2 (Fig. 3D and Fig. 4A-C). Since the actual measured redox state of each probe is affected similarly to natural redox-regulated proteins, by the combining activity of reductive and oxidative signals, the comparison between the oxidative states of these probes allows deducing 2-Cys Prxs oxidizing activity. This concept is clearly demonstrated in the response of the three biosensors to low light conditions (Fig. 4A-C). The immediate reduction observed for the chl-roGFP2 and roGFP2-Prx probes signifies the dominant reductive activity during the dark to low light transition. Most strikingly, the relatively stable oxidation state of chl-roGFP2-PrxΔC_R_ at the onset of low light compared to the reduction observed in the chl-roGFP2 and chl-roGFP2-Prx probes, at the same conditions, suggest the induction of oxidative signals under low-light intensities, attenuating the reductive signals emanated from GSH/GRXs, FTR/Trx and NTRC activities. These results are in agreement with the observations of transmitted oxidative signals through ACHT1/4 under low to moderate light intensity^23,24^.

Tracking the in vivo dynamics of oxidative thiol modulations is central to understanding the role of redox regulation in adjusting metabolic activity. Exploring the diurnal redox dynamics of the CBC enzymes, FBPase and SBPase, showed a change from fully oxidized to an almost fully reduced state of FBPase during gradual increases in light intensity from dark to 350 μmol photons m^−2^ s^−1 44^. Intriguingly, the observed light range in which FBPase reduction occurred matched the range of light-induced changes in 2-Cys redox signaling, as observed either by direct measurements of the 2-Cys Prx redox state or by using the chl-roGFP2-PrxΔC_R_ (Fig. 1E, Fig. 3D and Fig. 4A). Considering the sensitivity of FBPase to 2-Cys Prx-mediated oxidation^27^, these results suggest that the reductive and oxidative pathways are interconnected in regulating the activation state of carbon assimilation enzymes under low- and moderate-light conditions. This integrative view suggests that eliminating oxidative signals will result in a higher reduction state of CBC enzymes and consequently higher carbon assimilation rates. Indeed, the high reduction state of FBPase recently reported in *2cpab* plants^45^ and the rapid increase in carbon assimilation rates observed exclusively in *2cpab* and not in *prxQ* or *prxIIE* mutants plants (Fig. 4D) corroborate this view and points to the role of 2-Cys Prx in attenuating carbon assimilation rates. Furthermore, the inverse relationship between NADPH level and the induction of oxidative signals (Fig. 3D and Fig. 4A) suggests a feedback mechanism in which NADPH consumption may trigger oxidative signals, leading to inhibition of CBC enzyme activities and a consequent decrease in carbon assimilation rates.

In contrast to the rapid increase in carbon assimilation rates observed in *2cpab* plants during the photosynthesis induction phase, lower steady-state carbon assimilation rates under HL in *2cpab* plants compared to wild-type were recently reported^45^ and also observed in this study (Supplementary Fig. 7). Thus, 2-Cys Prx affects photosynthesis performance by two means. Under non-stress and conditions, 2-Cys Prx redox signaling, which is involved in attenuating carbon assimilation rates, is prevalent. On the contrary, under steady-state HL conditions, in which a higher rate of peroxide production is expected, the antioxidant activity rather than the signaling role of 2-Cys Prx is dominant and protects the photosynthetic apparatus from oxidative damage^16^.

Due to the heterogeneous organization and spatial arrangement of crop canopies, leaves lower in the canopy experience sporadic HL, scattered with low-light levels of variable durations. Given the dominant role of redox signaling at low to moderate light intensities, the level of activation and deactivation of CBC enzymes in lower leaves will vary throughout the day, with both the reductive and oxidizing signal shaping the rate of carbon assimilation rate, defining the contribution of these leaves to the total carbon assimilation rates^46^. The inherent redox changes that stimulate the activation and deactivation of CBC enzymes link carbon fixation to electron transport activity and energy flux but also govern limitation to optimal canopy assimilation under fluctuating light intensities. Given the role of 2-Cys Prx in deactivating CBC enzyme activity, our data suggest that the chl-roGFP2-PrxΔC_R_ probe estimates the degree of limitation of photosynthesis imposed by oxidative signals in plants, laying the foundation for increasing photosynthesis efficiency through redox modulations.

## Materials and Methods

### Plant material, growth conditions and treatment

*Arabidopsis thaliana* WT (ecotype Columbia), *prxQ* (Salk_070860C) and *prxIIE* (Salk_203706C) were obtained from ABRC. The mutants, *2cpab, ntrc* and *ntrc-Δ2CP* were obtained from Prof. Francisco Javier Cejudo^26,43^. WT and mutants plants were grown in a greenhouse under a 16/8 h light/darkness cycle. Immunoassay, redox imaging and thiol labelling experiments were performed on whole, three-week-old *Arabidopsis* plants. For the 3-(3,4-dichlorophenyl)-1,1-dimethylurea (DCMU) treatment, three-week-old plants were sprayed with 150μM DCMU (D2425-100G, Sigma) in the darkness phase, 1 hour before the onset of the light period.

### Construction of chl-roGFP2-PrxΔC_R_ and chl-PrxΔCR-roGFP2 probes

Both fusion probes (chl-roGFP2-PrxΔC_R_ and chl-PrxΔCR-roGFP2) were synthesized with codons optimized for *Arabidopsis* expression (Genewiz). Chloroplast targeting was achieved by using the 2-Cys peroxiredoxin A signal peptide. The genetically fused sequences were cloned into the plant cloning vector pART7, using the XhoI and HindIII restriction enzymes. The whole construct, including the CaMV 35S promoter and ocs terminator, was then cloned into the binary vector pART27, using the NotI restriction enzyme. The pART27 plasmids containing the chl-roGFP2-Prx, chl-roGFP2-PrxΔC_R_ and chl-PrxΔCR-roGFP2 probe constructs, were transformed into GV3101 Agrobacterium tumefaciens. Transformation of *Arabidopsis* thaliana (Columbia, Col-0) was then performed by floral dip^47^. Transformant lines were selected based on kanamycin resistance and the chl-roGFP2 fluorescence signal.

### Screening of chl-roGFP2-Prx and chl-roGFP2-PrxΔC_R_ DNA insertion in transformed plants

The substitution of the resolving cysteine (C_R_) by alanine in chl-roGFP2-PrxΔC_R_ plants was verified by DNA insertion screening. DNA extraction and PCR amplification were carried out using REDExtract-N-AmpTM Plant PCR kit (sigma-aldrich). The sequence of both primers, forward and reverse, are as follows: Forward primer: CATCCAAGAGAACCCAGATGAAGCT; Reverse primer: TTGCAGAGAAATATTCTTTAGACAAC. The Forward primer was designed to form HindIII restriction site sequence in the PCR products of Prx, thus allowing us to distinguish between Prx and PrxΔC_R_. PCR product for Prx (80bp):

> CATCCAAGAGAACCCAGATGAAGCTTGCCCAGCTGGTTGGAAACCCGGTGAGAAGTCTATGAAGCCCGATCCTAAGTTGTCTAAAGAATATTTCTCTGCAA

PCR product for PrxΔC_R_ (101bp):

> CATCCAAGAGAACCCAGATGAAGCTGCTCCAGCTGGTTGGAAACCCGGTGAGAAGTCTATGAAGCCCGATCCTAAGTTGTCTAAAGAATATTTCTCTGCAA

The PCR products were digested with HindIII and result in 80bp product for Prx and 101bp for PrxΔC_R_ (Supplementary Fig. 3).

### NEM-based redox labeling

Protein extracts were prepared in the presence of 1 ml cold 10% trichloroacetic acid (TCA) (dissolved in water) to retain the in vivo thiol oxidation status of proteins. Proteins were precipitated for 30 min on ice, in the darkness, and centrifuged at 19,000 xg for 20 min, at 4°C. The pellet was then washed four times with 100% cold acetone. After removal of the residual acetone, the pellet was resuspended in urea buffer (8M urea, 100mM 4-(2-hydroxyethyl)-1-piperazineethanesulfonic acid (HEPES) (pH 7.2), 1mM EDTA, 2% (w/v) SDS, protease inhibitors cocktail (PI) (Calbiochem) and 100mM N-ethylmaleimide (NEM) (E3876, Sigma)) dissolved in ethanol. Samples were then incubated for 30 min, at room temperature, followed by centrifugation (19,000 xg, 20 min, 4°C) and washed (4 times) with 100% cold acetone. The dry pellets were resuspended in urea buffer without NEM. The sample buffer (x3) contained 150mM Tris-HCl, pH 6.8, 6% (w/v) SDS, 30% glycerol and 0.3% pyronin Y. Gel fractionation was carried out without a reducing agent, on a precast 4–15% polyacrylamide gel. Fractionated proteins were transferred to a polyvinylidene fluoride membrane (Bio-Rad), using the Trans-Blot Turbo Transfer System (Bio-Rad) with Trans-Blot Turbo Midi Transfer Packs. The membrane was incubated with anti-PRX antibodies (1:1,000) (kindly provided by Prof. Avichai Danon) or an anti-peroxiredoxin-SO_3_ antibody (Abcam, ab16830, 1:2000), both followed by anti-rabbit horseradish peroxidase (HRP)-conjugated IgG (1:20,000) (Agrisera). Chemiluminescence was detected using the Advanced Molecular Imager HT (Spectral Ami-HT, Spectral Instruments Imaging, LLC., USA).

### mPEG-based redox labeling

Protein extract was prepared, using a hand homogenizer, in 1ml cold 10% trichloroacetic acid (TCA) (dissolved in water) to retain the in vivo thiol oxidation status of proteins. Proteins were precipitated for 30 min, on ice, in the darkness, and then centrifuged at 19,000 xg for 20 min at 4°C. The pellet was washed four times with 100% cold acetone, after which, residual acetone was removed and the pellet was resuspended in urea buffer with NEM. The samples were then incubated for 30 min at room temperature. Samples were then reduced by addition of 100mM DTT (60 min, room temperature). TCA (10%) was then added and samples were precipitated for 30 min, on ice, in the darkness. The TCA-treated extracts were then centrifuged at 19,000 xg, 4°C for 20 min, supernatants were removed and the pellets were washed three times with 100% acetone. The dry pellets were resuspended in urea buffer containing 10mM methoxy-polyethylene glycol maleimide (mPEG, MW 5000 g/mol, Sigma), and incubated at 28 °C for 2 h. The reaction was stopped by adding an equal volume of sample buffer containing 50 mM DTT. Samples were then separated by 10% Tricine–SDS–PAGE and then transferred to a polyvinylidene fluoride membrane (Bio-Rad) under the same conditions described above. The membrane was incubated with anti-PRX antibodies (1:1,000) and then with anti-rabbit HRP-conjugated IgG (1:20,000).

### Probe detection using western analysis

Experiments were performed on 10-day-old seedlings. Whole-plant contents were extracted with extraction buffer (20 mM Tris, pH 8.0, 1 mM EDTA, and 50 mM NaCl), and then centrifuged at 19,000 xg, for 20 min, at 4°C. Gel fractionation of 100 μg extracted protein was carried out by reducing 10% SDS-PAGE. Fractionated proteins were then transferred onto a polyvinylidene fluoride membrane, which was then incubated with anti-PRX antibodies (1:1,000) or anti-GFP (abcam) antibodies (1:5,000), and then with anti-rabbit HRP-conjugated IgG (1:20,000).

### Confocal microscopy

Confocal microscopy analysis was carried out on 2-week-old transgenic chl-roGFP2, chl-roGFP2-Prx, chl-roGFP2-PrxΔC_R_ and chl-PrxΔCR-roGFP2-expressing plants. Images were captured with a Leica TCS SP8 confocal system (Leica Microsystems) and the LAS X Life Science Software, while using a HC PL APO ×40/1.10 objective. To capture chl-roGFP2 fluorescence, samples were excited at 488 nm and emission was measured at 500–520 nm. For chlorophyll fluorescence, excitation was at 488 nm and emission was at 670 nm. Merged images were generated using Fiji (Image 1.A) software.

### roGFP2 fluorescence measurements and analysis

Redox imaging was performed on three-week-old transgenic chl-roGFP2 and chl-roGFP2-PrxΔC_R_-expressing plants. Whole-plant roGFP2 fluorescence was detected using an Advanced Molecular Imager HT, and images were taken using the AMIview software. For roGFP fluorescence images, excitation was at 465nm (reduction) and 405nm (oxidation), followed by emission at 510nm. Chlorophyll fluorescence, which was used to select plant tissue pixels, was measured following excitation at 405 and emission at 670. For each probe, images were taken under the same settings of LED intensities and exposure time. Ratiometric images were created by dividing, pixel by pixel, the 405 nm image by the 465 nm image, and displaying in false colors. Images were processed using a custom-written Matlab script. For calibration of the probe response, detached plants were immersed in 0.5M H_2_O_2_ or 100 mM DTT, and ratiometric images for fully oxidized and fully reduced states, respectively, were obtained.

Daily roGFP2 fluorescence was measured using a Tecan Spark^®^ multimode microplate reader^48^. For all probes (chl-roGFP2, chl-roGFP2-Prx and chl-roGFP2-PrxΔC_R_), roGFP fluorescence was measured using excitation at 485nm/20 (reduction) and 400nm/20 (oxidation), followed by emission at 520nm/10; gain values were adjusted to avoid signal saturation. For chlorophyll detection, 400nm/20 was applied for excitation, followed by 670nm/40 for emission. For automatic detection of chl-roGFP2 signals, 10-day-old plants expressing the roGFP2-based probes or wild type plants were transferred to solid peat plugs in 12-well plates. The plates were placed in a growth chamber (Fytoscope FS-SI-4600, Photon Systems Instruments, Czech Republic) under normal growth conditions for another 5 days before the start of the experiment. An automated robot (KiNEDx KX.01467, paa automations, Hampshire, UK) controlled by a self-compiled program (Overlord™, paa automations), inserted the plates into the plate reader to measure chl-roGFP2 fluorescence. For the plate-reader analysis, a 9-by-9-pixel matrix was formed for each well. Chlorophyll fluorescence was detected to create a chlorophyll mask, which was then used to choose pixels that returned a positive chlorophyll fluorescence signal, and only those pixels were considered for the roGFP analysis. The average signals of non-fluorescent plants (WT) was calculated, and the values were deducted from the values detected in the chl-roGFP2 fluorescence analysis. A similar experimental setup was used to measure the fluorescence emitted from the TKTP-iNAP4 and TKTP-iNAPc expressing lines. The pH corrected ratio for TKTP-iNAP4 was calculated by dividing the fluorescence ratio values obtained from the TKTP-iNAP4 line by those obtained from the TKTP-iNAPc line, according to^39^. To determine sensor OxD, detached roGFP-based probe-expressing and WT plants were immersed in 0.5M H_2_O_2_ or 100 mM DTT and fluorescence was measured. roGFP2 OxD was calculated based on the fluorescence signal according to Equation^31^.

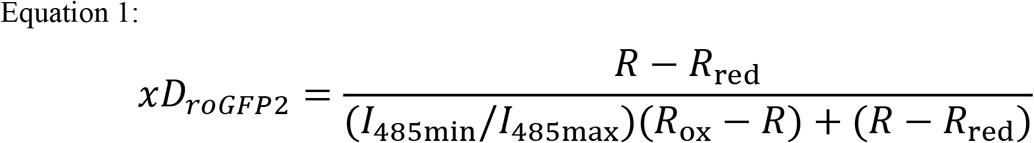

where R represents the ratio (400/485) at each point in the experiment, Rred represents the ratio under fully reduced conditions, Rox represents the ratio under fully oxidized conditions, I485min represents the fluorescence emitted at 520 nm when excited at 485 nm under fully oxidized conditions and I485max represents the fluorescence emitted at 520 nm when excited at 485 nm under fully reduced conditions.

A self-written Matlab script was used to analyze the many files that contained the measurements of each experiment. In addition, experimental metadata such as plant lines, light irradiance, temperature, relative humidity and CO_2_ concentration were also collected and added to the output file.

### Gas Exchange Measurements

A gas exchange measurements were performed using the LI-6800 (LI-COR^®^). Four plants were planted in LI-COR^®^ 65mm pots (610-09646) and analysis was done using the LI-6800 small plant chamber (6800-17). Total leaf area was calculated via image analysis using the Ami HT Imager and Matlab.

## Supporting information

Supplementary Figuers

## Author contributions

N.L. and S.R. conceived and designed the research. N.L., I.N., R.L., G.G., M.H., and S.R. performed the research. N.L., R.L., M.H., and S.R. analyzed the data. N.L. and S.R. wrote the manuscript. Competing interests: The authors declare no competing interests.

## Acknowledgments

We thank Avihai Danon for his critical comments on the manuscript and for kindly providing us the anti-2-Cys Prx antibody. We thank Francisco Javier Cejudo for kindly providing *2cpab, ntrc* and *ntrc-Δ2CP* mutants. We thank Wallace Boon Leong Lim, who kindly provided the TKTP-iNAP4 and TKTP-iNAPc Arabidopsis lines. We thank Bruce Morgan for his kindly help in initiating this project.

## Funding

This research was supported by the Israel Science Foundation (grant No.826/17 and No. 827/17).

## Notes

### Competing Interest Statement

The authors have declared no competing interest.

