## Supplementary Figuers for "Systematic monitoring of 2-Cys peroxiredoxin-derived redox signals unveiled its role in attenuating carbon assimilation rate"

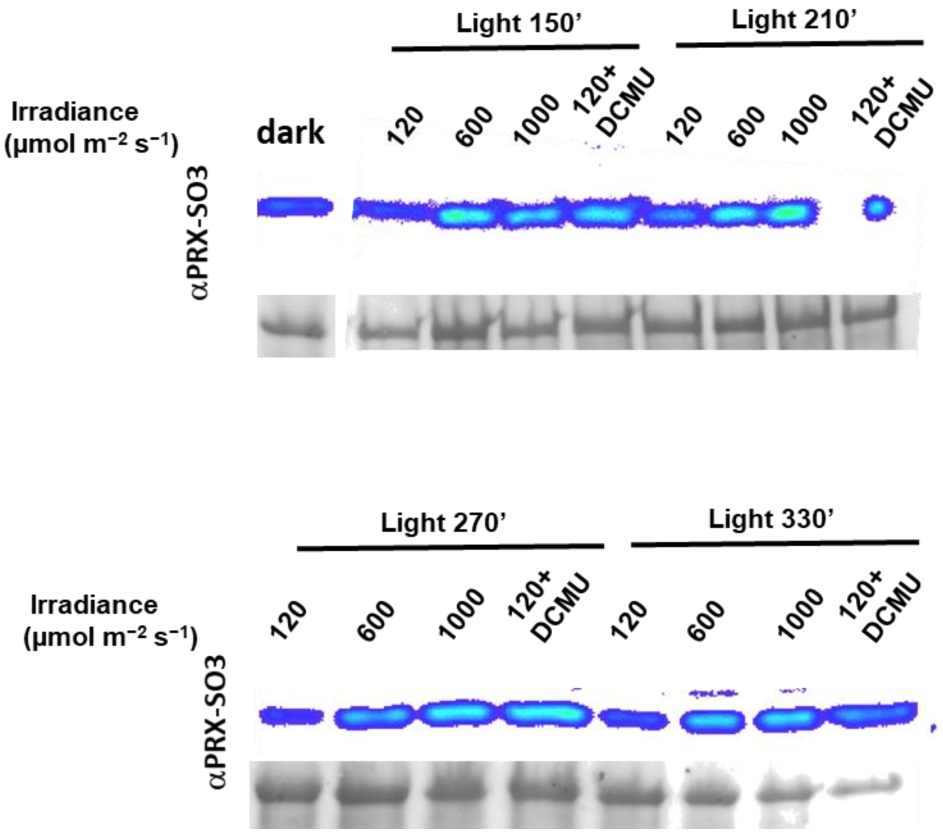


**Supplementary Figure 1. Hyperoxidation of 2-Cys Prx under various light conditions.** Immunoblot assay showing the hyperoxidiezed form, $SO_3$ in plants exposed to prolong high-light conditions. Supports Fig. 1A.

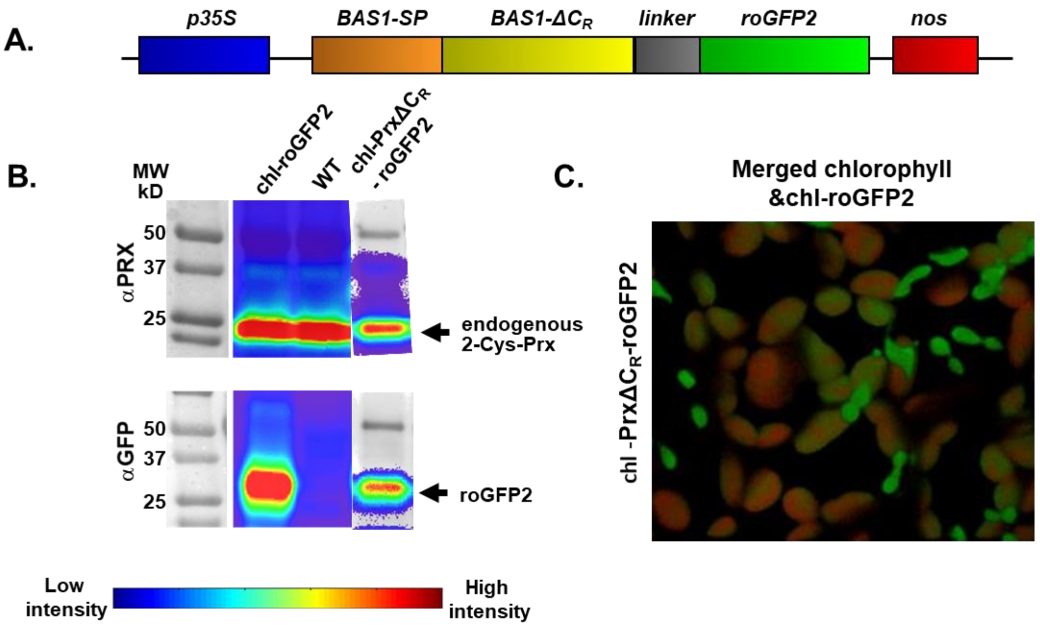


**Supplementary Figure 2. BAS1 is prone to cleavage by specific chloroplast proteases. (A)** Schematic diagram of the gene cassette used to transform Arabidopsis plants. **(B)** *In vivo* probe detection by western blot analysis using either anti-Prx or anti-GFP antibodies. **(C)** Subcellular localization of Prx$\Delta C_R$-roGFP2- in mesophyll cells, as detected by confocal microscopy. Supports Fig. 2, demonstrating the cleavage of Prx$\Delta C_R$-roGFP2 probe.

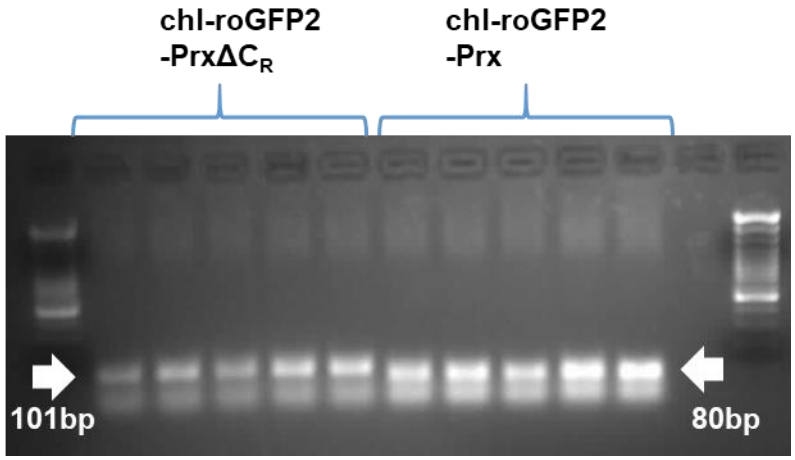


**Supplementary Figure 3. A restriction-based assay for the screening of chl-roGFP2-Prx and chl-roGFP2-PrxΔC$_R$ DNA insertions in transformed plants.** The substitution of the resolving cysteine (C$_R$) by alanine in chl-roGFP2-PrxΔC$_R$ plants was verified by DNA insertion screening (See Methods).

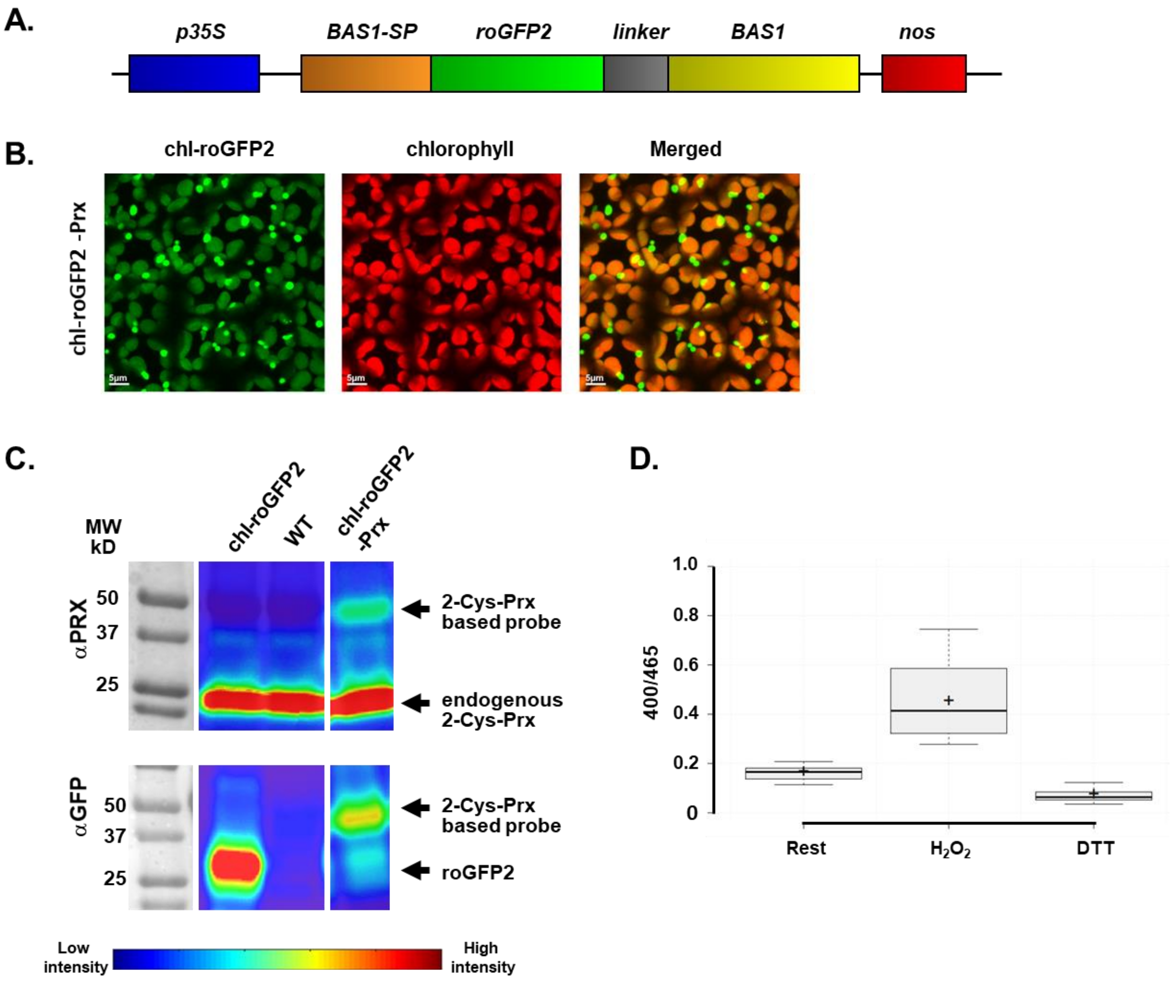


**Supplementary Figure 4. Chloroplast-targeted roGFP2-Prx probe. (A)** Schematic diagram of the gene cassette used to transform Arabidopsis plants. **(B)** Subcellular localization of chl-roGFP2-Prx in mesophyll cells, as detected by confocal microscopy. **(C)** In vivo detection of the fused sensor protein by western blot analysis using either anti-Prx or anti-GFP antibodies. **(D)** Fluorescence ratio (405/465) in plants expressing chl-roGFP2-Prx under steady-state conditions and following treatment with 100mM DTT or 500mM $H_2O_2$. The emitted fluorescence from three-week-old plants grown in soil was recorded using a plate reader.

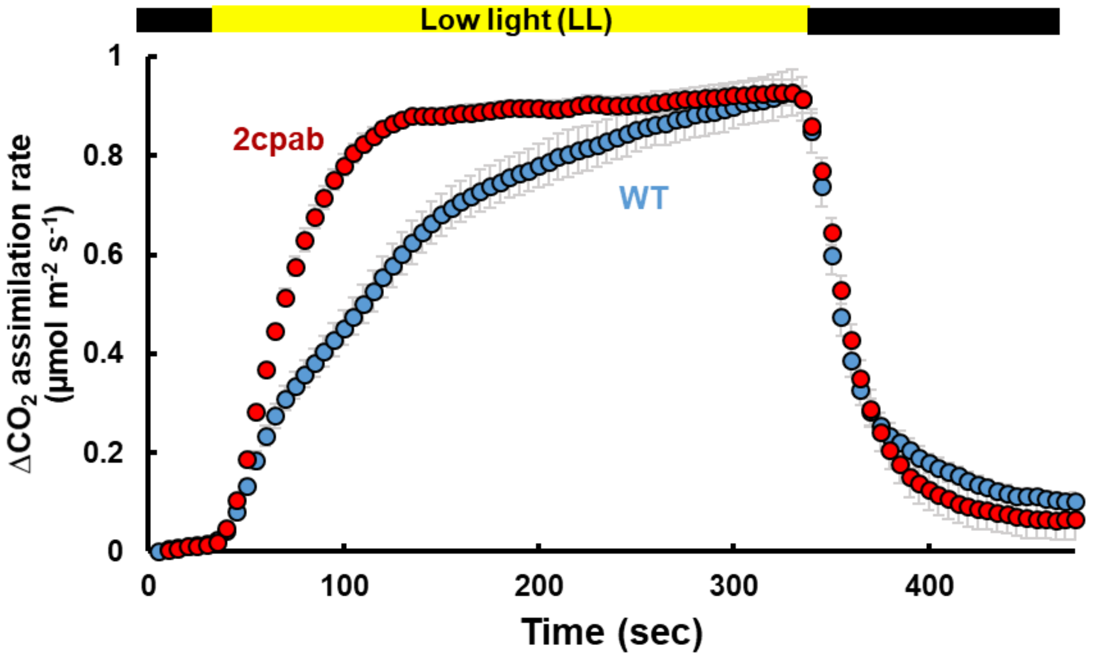


**Supplementary Figure 5**. **Carbon assimilation measurements upon dark to low light transition in *2cpab* and wild-type plant.** Carbon assimilation rates were measured in dark-adapted wild-type and *2cpab* mutants plants upon exposure to transient low light (5min of 22 µmol photons $m^{-2}$ $s^{-1}$). Data were normalized to values obtained during the first dark period. Values for the carbon assimilation are presented as means of 4-5 pots with 4 plants in each ± SE. The color bar denotes the light conditions: black-dark, yellow-light. Supports Fig.4.

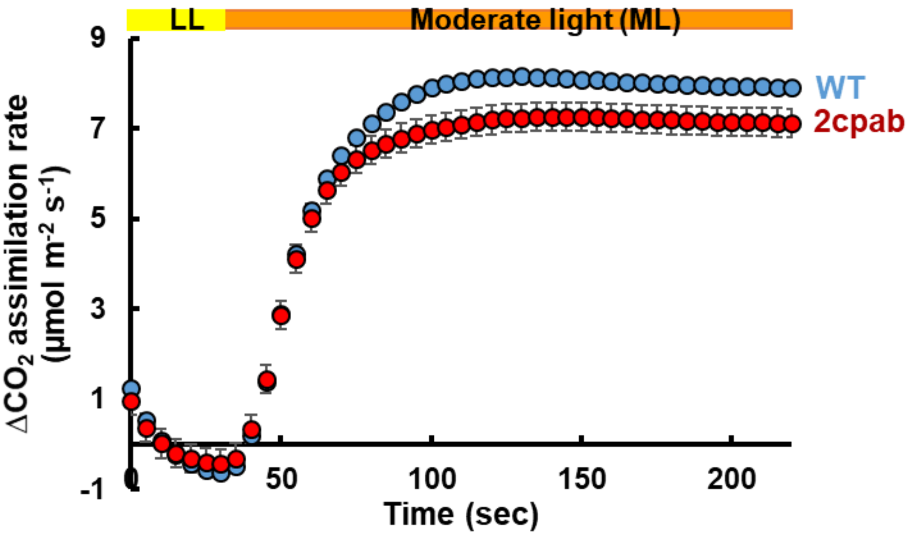


**Supplementary Figure 6. Carbon assimilation measurements under fluctuating light in *2cpab* and wild-type plants.** Carbon assimilation rates were measured in moderate light (330 µmol photons $m^{-2}$ $s^{-1}$) adapted WT and *2cpab* plants which were exposed to 1min of low light (22 µmol photons $m^{-2}$ $s^{-1}$) followed by moderate light (330 µmol photons $m^{-2}$ $s^{-1}$). Data were normalized to values obtained during low light at the beginning of the experiment. Values for the carbon assimilation are presented as means of 4 pots with 4 plants in each ± SE. The color bar denotes the light conditions: yellow- low light, orange-moderate light. Supports Fig.4.

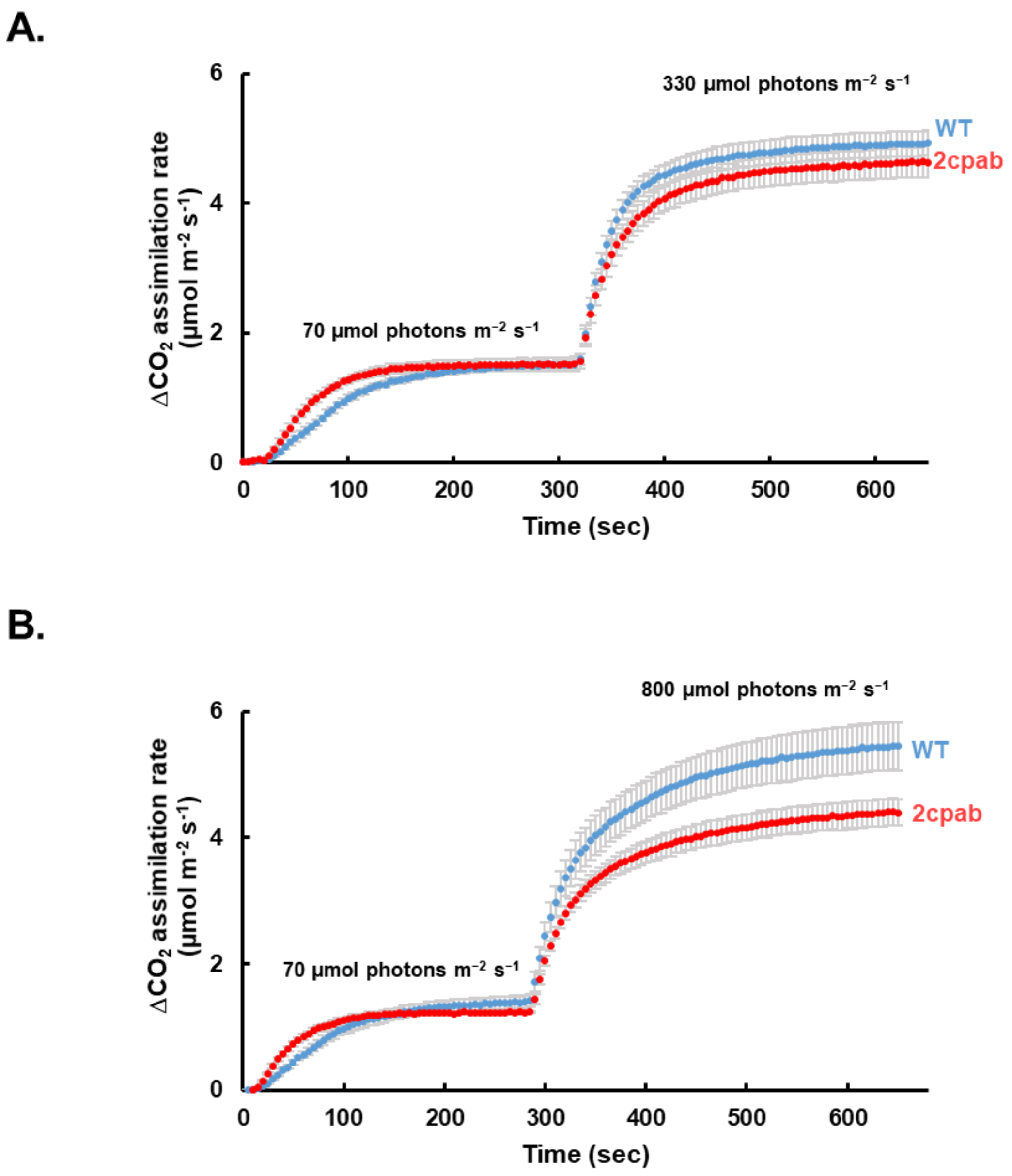


**Supplementary Figure 7. Carbon assimilation measurements under moderate and high light in *2cpab* and wild-type plants.** Carbon assimilation rates were measured in dark-adapted WT and *2cpab* mutants plants upon exposure to dark, followed by illumination of 70 µmol photons $m^{-2}$ $s^{-1}$ (5min) and transition to **(A)** 300 or **(B)** 800 µmol photons $m^{-2}$ $s^{-1}$. Data were normalized to values obtained during the first dark period. Values for the carbon assimilation are presented as means of 4-5 pots with 4 plants in each ± SE. Supports Fig.4.
